# Single-cell genomics reveals the divergent mitochondrial genomes of Retaria (Foraminifera and Radiolaria)

**DOI:** 10.1101/2023.02.03.527036

**Authors:** Jan-Niklas Macher, Nicole L. Coots, Yu-Ping Poh, Elsa B. Girard, Anouk Langerak, Sergio A. Muñoz-Gómez, Savar D. Sinha, Dagmar Jirsová, Rutger Vos, Richard Wissels, Gillian H. Gile, Willem Renema, Jeremy G. Wideman

**Author notes:** Shared first authorship (Jan-Niklas Macher, Nicole L. Coots).

## Abstract

Mitochondria originated from an ancient bacterial endosymbiont that underwent reductive evolution by gene loss and endosymbiont gene transfer to the nuclear genome. The diversity of mitochondrial genomes published to date has revealed that gene loss and transfer processes are ongoing in many lineages. Most well-studied eukaryotic lineages are represented in mitochondrial genome databases, except for the superphylum Retaria—the lineage comprising Foraminifera and Radiolaria. Using single-cell approaches, we present two complete mitochondrial genomes of Foraminifera and two near-complete mitochondrial genomes of radiolarians. We report the complete coding content of an additional 14 foram species. We show that foraminiferan and radiolarian mitochondrial genomes encode a nearly fully overlapping but reduced mitochondrial gene complement compared to other sequenced rhizarians. In contrast to animals and fungi, many protists encode a diverse set of proteins on their mitochondrial genomes, including several ribosomal genes; however, some aerobic eukaryotic lineages (euglenids, myzozoans, and chlamydomonas-like algae) have reduced mitochondrial gene content and lack all ribosomal genes. Similar to these reduced outliers, we show that retarian mitochondrial genomes lack ribosomal protein and tRNA genes, contain truncated and divergent small and large rRNA genes, and encode only 14-15 protein-coding genes, including nad1, 3, 4, 4L, 5, 7, cob, cox1, 2, 3, atp1, 6, and 9, with forams and radiolarians additionally encoding nad2 and nad6, respectively. In radiolarian mitogenomes, a non-canonical genetic code was identified in which all three stop codons encode amino acids. Collectively, these results add to our understanding of mitochondrial genome evolution and fill in one of the last major gaps in mitochondrial sequence databases.

## Introduction

Endosymbiosis, the functional integration of one cell into another, has shaped the evolution of eukaryotes (1, 2). The oldest extant endosymbioses, mitochondria and chloroplasts, both originate from ancient bacterial endosymbionts (3, 4). From their origin to the present, mitochondrial and chloroplast genomes have undergone reductive evolution by gene loss or endosymbiont gene transfer (EGT) to the nuclear genome (3, 5–7). For mitochondrial genomes, most of this reduction occurred prior to the emergence of the last eukaryotic common ancestor (LECA). What was once a bacterial genome encoding thousands of proteins became a mitochondrial genome encoding less than a hundred proteins in LECA (3). Plants and many protist lineages still retain a diverse set of between 30 and 70 protein-coding genes on their mitochondrial genomes (3, 8–11). In addition to electron transport chain (ETC) components from Complexes I-V, mitochondrial genomes can encode upwards of 40 other proteins involved in transcription, translation, complex maturation, and transport (9, 12). In extreme cases of reduction, the highly reduced mitochondria-related organelles (MROs) have entirely lost their organellar genomes (3, 13–15). Although these extreme cases of reduction are associated with anaerobic lifestyles, several aerobic lineages have also undergone major reductions in their mitochondrial genome content, mostly via EGT to the nucleus (16–19). Why the organelle retains certain mitochondrial genes is hotly debated (20–25), and it remains unclear what functional consequences varying gene complements might endow.

Mitochondria are best known for their role in synthesizing ATP using a proton gradient across their inner membrane (26). In fact, mitochondrial genomes largely code for proteins directly or indirectly related to the function of the ETC and ATP synthase (27). Furthermore, when the need for the ETC is lost in anaerobic lineages, the mitochondrial genome is also lost (14, 28). In aerobic lineages, a few components of the ETC (e.g., parts of Complex I, III, and IV) and the mitochondrial ribosomal RNAs are always encoded in mitochondrial genomes (29). Apart from these few components, all other genes can be lost, replaced (30), or transferred to the nuclear genome. In the lineages leading to animals and fungi, all genes encoding ribosomal proteins (except *rps3* in most fungi (31)) were independently transferred to the nuclear genomes (19). Thus, most animal and fungal mitochondrial genomes encode only rRNAs, tRNAs, and 13-14 proteins (fewer if Complex I is lost like in *Saccharomyces cerevisiae*), all of which are involved in electron transport or ATP synthesis (32).

In addition to the mitochondrial genome reductions seen in animals and fungi, some aerobic protist lineages also exhibit ancient reductions of their mitochondrial coding repertoire and contain many fewer proteins, few or no tRNAs, and highly divergent or fragmented rRNAs. For example, myzozoans, which include apicomplexans and dinoflagellates, encode only one to four ETC proteins in addition to highly fragmented and extremely divergent rRNAs (33–37); euglenids like *Euglena gracilis* encode 8 ETC proteins and very short divergent rRNAs (38, 39); and chlorophycean algae like *Chlamydomonas reinhardtii* encode 7 ETC proteins and extremely fragmented rRNAs (40, 41). In addition to these major aerobic lineages, one aerobic genus, the red alga *Galdieria* (*42*), has also lost its mitoribosomal proteins from its mitochondrial genomes via EGT. Thus, although mitochondrial genomes often encode many proteins, certain evolutionary circumstances result in massive gene loss or EGT, resulting in reduced mitochondrial genome coding content.

While several orphan taxa still lack representation in mitochondrial genome databases, only one major eukaryotic lineage is completely absent - the Retaria, the rhizarian lineage comprising the phyla Foraminifera and Radiolaria (43). As a whole, rhizarians are important members of marine communities (44–47), contributing significantly to marine biogeochemical cycling (48–50). Retarians are aerobes (although some forams thrive under anoxic conditions (51)) and are abundant in many environments, especially in the global ocean. With ~9000 recognized mostly marine extant species, Foraminifera are estimated to account for ~25% of present-day carbonate production (52, 53). Silicified Radiolaria, with their 600-800 named species, are estimated to account for between 2-19% of total biogenic silica production (54). Despite their importance, the paucity of retarian genomes and transcriptomes in sequence databases has made a deeper understanding of these lineages impossible (55–60).

In order to obtain mitochondrial genome sequences from Retaria, we chose to use single-cell approaches. Single-cell genomics can effectively recover mitochondrial genomes from diverse protists (10, 61, 62). Even though most species of Foraminifera and Radiolaria are not in culture, contain a multitude of symbionts (63, 64), and show high levels of intragenomic polymorphisms (65, 66), we show that single-cell approaches can effectively recover mitochondrial genomes from these complex assemblages. Our data demonstrate that foraminiferan and radiolarian mitochondrial genomes encode an overlapping but reduced gene complement compared to other sequenced rhizarians, similar to other reduced mitochondrial genomes from other lineages. Retarian mitochondrial genomes do not encode ribosomal proteins or tRNAs. However, they do contain truncated and divergent small and large rRNAs and encode only 14-15 protein-coding genes, including *nad1, 3, 4, 4L, 5, 7, cob, cox1, 2, 3, atp1, 6,* and *9*, with forams and radiolarians additionally encoding *nad2* and *nad6*, respectively. An alternative genetic code was identified in radiolarian mitogenomes in which all three stop codons encode amino acids (TGA = W, TAG = Y, and TAA = Y/STOP). These results further add to our understanding of mitochondrial genome evolution across the eukaryotic tree of life.

## Results and Discussion

### Retarian mitochondrial, but not nuclear genomes, can be readily recovered using single-cell methods

We isolated individual cells, and Illumina-sequenced and assembled mini-metagenomes of 31 Foraminifera from 15 species (which are impossible to separate from their symbionts) and single-cell amplified genomes (SAGs) of 5 Radiolaria from 2 species (see Table S1 for a complete list). Foram mini-metagenomes are referred to as SAGs henceforth. One additional foraminiferan metagenome (*Globobulimina sp.*) was downloaded and reassembled from the NCBI sequence read archive (accession number SRA SRX3312059 (67)). Assemblies from *Calcarina*, *Neorotalia*, *Lithomelissa*, and *Acanthometra* SAGs are available to BLAST at SAGdb (https://evocellbio.com/SAGdb/macher_et_al/).

Both forams and radiolarians associate with many eukaryotic and bacterial endosymbionts (68–70), making it difficult to obtain *bona fide* sequence data from either lineage. To assess the contamination in foram and radiolarian SAGs, we collected all 18S and 16S sequences from all assemblies using *Cafeteria roenbergensis* 18S and *E. coli* 16S sequences as BLAST queries. We found foraminifera 18S genes or gene fragments in 23 of 31 SAGs from 13 of 15 species (Table S1). We also identified specific symbiont 28S (from dinoflagellate symbiont) or rbcL (from diatom symbionts) in all foram SAGs and species except *Calcarina mayori* and the reassembled *Globobulimina* (which is expected, since *Globobulimina* does not contain photosymbionts) (Table S2). The inability to identify the foram 18S genes in all specimens is likely due to their extreme within-cell variability (65, 71), which prevented proper assembly. 16S BLAST searches recovered diatom symbiont chloroplast and mitochondrial genes. In addition, 16S sequences from two common bacterial genera were also recovered (*Burkholderia* and *Cutibacterium*). Blobplots from foram assemblies confirm 18S BLAST findings as large contigs of symbiont organelles (See Figure S1A-B). From these data, we concluded that our foram SAG assemblies predominantly contain symbiont contigs, with only some representation from the host nuclear genome.

In radiolarians, we obtained high-coverage contigs with complete 18S sequences only from radiolarians (Table S2). In *Acanthometra* and *Amphibelone* (nc69, 78, 87, 96) SAGs, a few contaminating 18S sequences (e.g., diatom, cryptophyte, ciliate) were detected, but these contigs were fragmented with low coverage indicating relatively few eukaryotic contaminants. Similarly, only a few fragmented low-coverage 16S contigs were recovered, again indicating very little prokaryotic contamination. These results are corroborated by blobplots showing relatively little contamination in Acantharian SAGs (Figure S1C). In the *Lithomelissa* SAG (r2m), only radiolarian 18S sequences were recovered. However, many high-coverage bacterial contigs containing 16S sequences were identified, indicating that eukaryotic contamination in this SAG was low, but bacterial contamination was very high. These 18S and 16S results are corroborated by blobplots showing a degree of bacterial and eukaryotic contamination, but a large proportion of ‘unknown’ reads with no similar hits in the NCBI non-redundant database (Figure S1D). To assess the contamination of our nuclear data, we used a phylogenetic placement approach to assess SAG contamination (Figure S2). Forams were excluded because they lacked sufficient identifiable nuclear contigs. Briefly, we extracted BUSCO proteins from SAG assemblies and added them to existing alignments from EukProt (72). Even with low radiolarian BUSCO scores (nc69 10.6%, nc78 5.1%, nc96 6.3%, for *Acanthometra,* nc87 3.9% for *Amphibelone*, and 11.4% for *Lithomelissa* SAGs), *Acanthometra* SAGs were correctly placed with full support alongside the only radiolarian (*Astrolonche serrata*) in the EukProt dataset (Figure S2). Conversely, the *Lithomelissa* SAG was placed within alveolates with full support (Figure S2), suggesting unseen eukaryotic contamination—even though no contaminating 18S could be detected. Collectively, these data indicate that our radiolarian SAGs contain a substantial amount of radiolarian nuclear contigs, though the *Lithomelissa* SAG contains a large degree of bacterial and possibly eukaryote contamination.

Since mitochondrial genomes are often overrepresented in genome assemblies (10), we sought to identify foram and radiolarian mitochondrial genomes in our SAGs. Using protein sequences encoded by the *Andalucia godoyi* mitochondrial genome, one of the most gene-rich mitogenomes known (12), we identified several putative retarian mitochondrial contigs in foram and most radiolarian SAGs. *Amphibelone* SAG nc87 (96% identical 18S to other nc SAGs) lacked any obvious mitochondrial contigs and was not investigated further. Since many assemblies were contaminated by both eukaryotic and prokaryotic contamination, great care was taken to inspect the validity of each contig manually. In forams, the putative mitochondrial contigs had orders of magnitude higher read coverage and much lower GC content than symbiont or putative foram nuclear contigs (seen clearly in bobplots Figure S1A-B). Contigs representing near complete or complete symbiont organelle genomes were also found in many foram SAGs, though these contigs had much lower coverage than the foram mitochondrial genomes (Figure S1A-B). In radiolarians, the results were less clear-cut. While the coverage (~40-60x) of the putative mitochondrial contigs was much higher than the median of the SAG (~3-5x for the *Acanthometra* SAGs and ~11 for *Lithomelissa* - likely due to some very high-coverage contigs - see Figure S3), the GC content was similar to that of the putative nuclear contigs (Figure S1). Thus, though these contigs had relatively high coverage, they were not clearly separated from the majority of contigs in blobplots. The coverage of mitochondrial genome contigs and contigs containing radiolarian 18S sequences had similar coverage (~ 40-60x). Since both mitochondrial genomes and 18S sequences are generally found in multiple copies in a cell, we reasoned we have likely sequenced *bona fide* mitochondrial genomes and not nuclear mitochondrial genomes (NuMts), which would likely have much lower read coverage.

### Retarian mitochondrial genomes encode a reduced gene complement

From each assembly, we extracted mitochondrial contigs collectively representing the putatively complete mitochondrial gene complement from 16 foraminiferan and two radiolarian species. We recovered contigs with mitochondrial genes from two species of radiolarians (*Lithomelissa* sp. and *Acanthometra* sp.) that almost completely overlap the foraminiferan complement (Figure 1).

**Figure 1.**
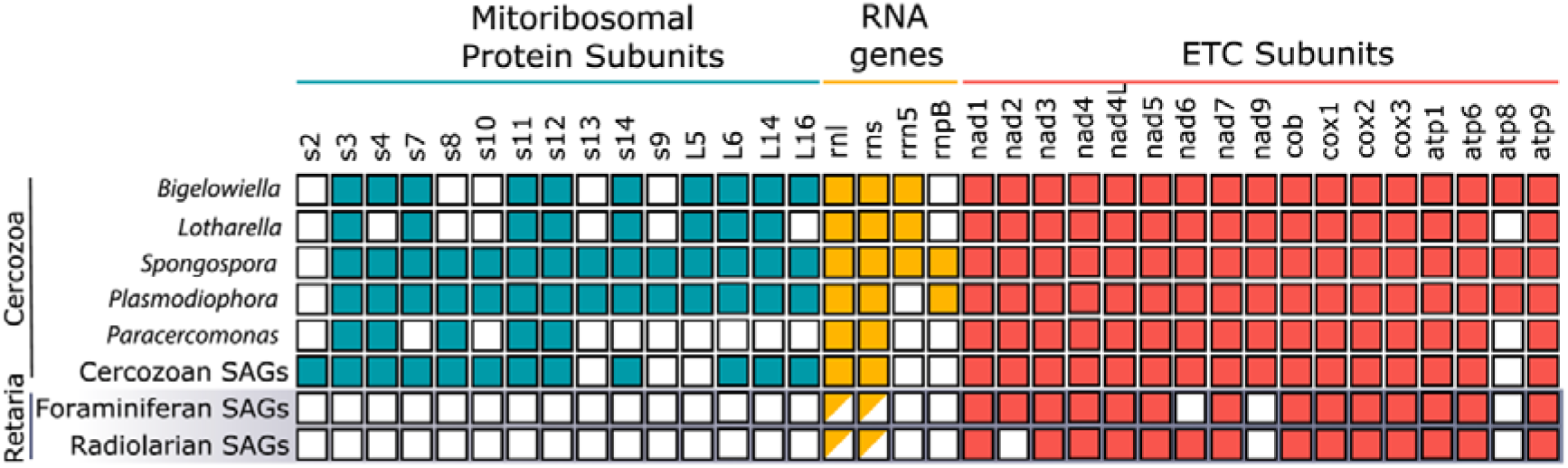
Mitochondrial genomes of forams and radiolarians overlap in gene content. Gene content of rhizarian mitochondrial genomes. Half-filled boxes indicate the presence of fragmented and shortened genes. Mitochondrial genes present in other mitochondrial genomes but absent from all sequenced rhizarian mitochondrial genomes are not listed. ‘Cercozoan SAGs’ refers to the single-amplified genomes published previously (10).

We obtained complete circular-mapping mitochondrial genomes of the forams *Calcarina hispida* and *Neorotalia gaimardi*. The mitogenomes were 46kb (*Calcarina hispida*) and 50kb (*Neorotalia gaimardi*) long, and each encoded the same set of 14 protein-coding genes (Figure 2). We concluded from these data that we likely extracted the complete, or near-complete, coding complement of these radiolarian mitochondrial genomes. However, we were unable to recover complete circular mitochondrial genomes from either radiolarian, likely due to repetitive intergenic regions that prevented proper assembly. To complete these genomes, we attempted to link contigs together using primers designed to PCR amplify missing regions between contigs but were unsuccessful—likely due to complex repetitive regions.

**Figure 2.**
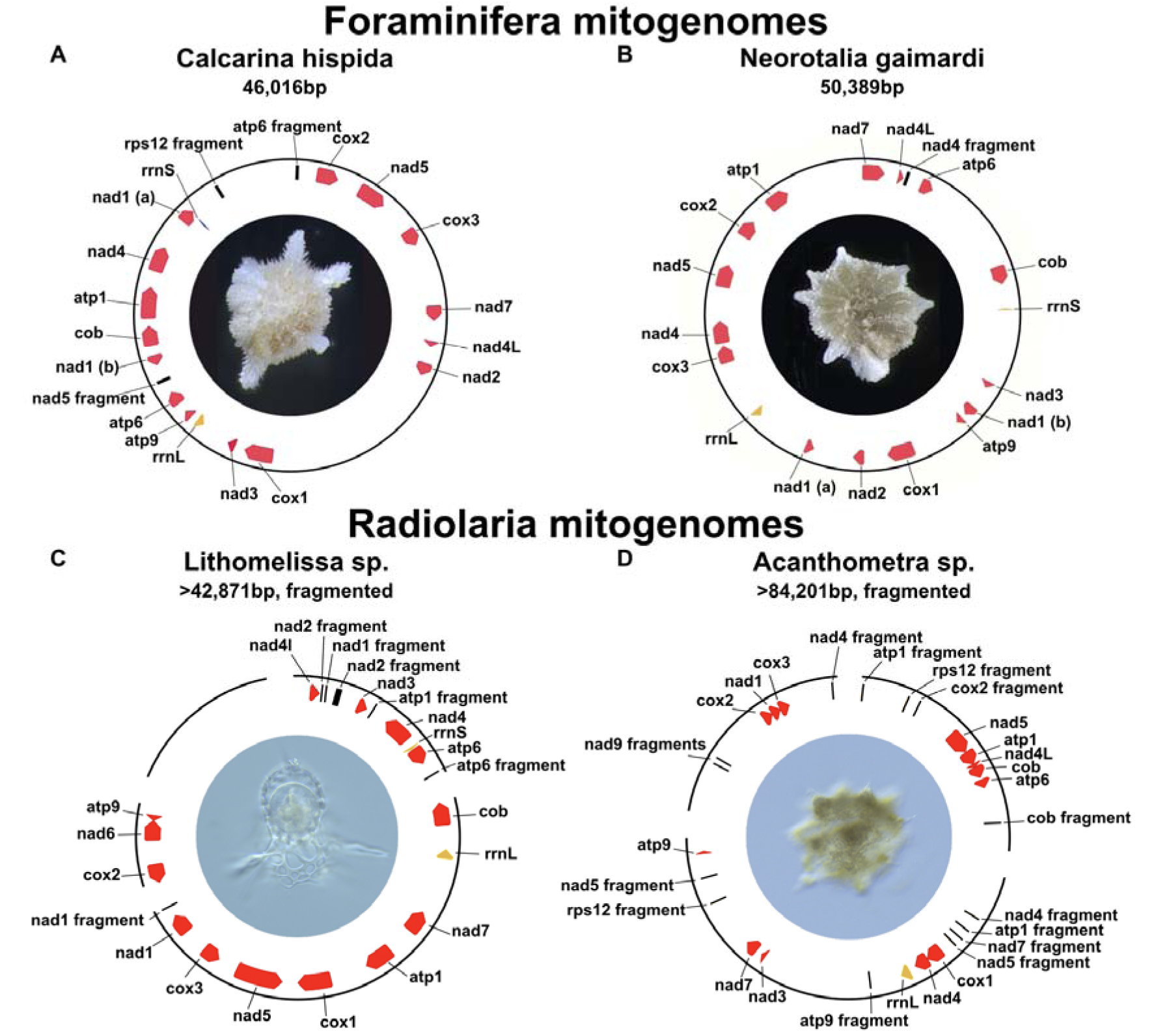
Retarian mitochondrial genomes contain large non-coding intergenic regions. Complete circular-mapping mitochondrial genomes of the Foraminifera *Calcarina hispida* (A) and *Neorotalia gaimardi* (B) and the inferred near-complete mitogenomes of the Radiolaria *Lithomelissa* sp. (C) and *Acanthometra* sp. (D). Protein-coding genes are highlighted in red, rRNAs (rrnL and rrnS) are highlighted in yellow. Gene fragments are shown in black. Gaps in the radiolarian mitochondrial genomes show the start and end of assembled mitochondrial contigs. Photos were taken of each individual organism before lysis.

Between the two phyla, retarians displayed a near-identical gene complement, including cytochrome *c* oxidase subunits (*cox1*, *cox2*, *cox3*), cytochrome *b* (*cob*), and ATP synthase subunits (*atp1*, *atp6*, and *atp9*), and NADH dehydrogenase subunits (*nad1*, *nad3*, *nad4*, *nad4L*, *nad5*, nad7; *nad2* is missing from radiolarians and *nad6* is missing from foraminifera), The lack of *nad2* in both radiolarian mitochondrial genomes and the lack of *nad6* in one radiolarian and both foram mitochondrial genomes is not without precedent as both are either lost or extremely diverged and transferred to the nuclear genome in euglenids (73). Fragments of LSU rRNA (*rrnL*) and SSU rRNA (*rrnS*) genes were identified in the mitogenomes of the foraminiferans *Calcarina hispida, Neorotalia gaimardi*, and the radiolarian *Lithomelissa sp.*, while only *rrnL* was identified in the mitogenome of the radiolarian *Acanthometra sp.*; however, neither full-length ribosomal protein-coding genes nor tRNAs were detected (Figure 1). The *nad9* gene was not found in our retarian mitochondrial genomes even though all other sequenced rhizarian mtDNAs contain this gene (10, 74–77) (except for *Brevimastigomonas*, which has lost Complex I altogether). Since most core Complex I subunit genes appear to be retained in rhizarian mitogenomes (including those of retarians), missing Complex I genes could be encoded by the nuclear genome; however, these genes have not been identified in the nuclear genomes of euglenids (73). However, we were unable to find any Complex I components in the nuclear assemblies likely indicative of their incompleteness (BUSCO scores <10%). Another conspicuous absence from retarian mitochondrial genomes is *atp8*, which encodes for subunit 8 of ATP synthase. Subunit 8 is likely an essential component of ATP synthase in most organisms (77) but appears to be absent in *Caenorhabditis elegans* (78) and cannot be identified in many rhizarian mitochondrial genomes (77, 79). To further confirm that we have collected *bona fide* mitochondrial contigs, we reconstructed the phylogeny of forams with radiolarians as an outgroup using concatenated mitochondrial proteins predicted from the contigs (Figure S4). The resulting phylogeny at the family level recapitulates the topology seen in 18S rDNA trees of Foraminifera (80–82), except for Peneroplidae clustering within the Soritidae.

We also obtained 25 fragmented mitochondrial genomes from 14 additional foraminiferan species (see Table S1 for a list of samples) that could not be linked in a single contig but had the same set of 14 protein-coding genes (all for ETC subunits) present in the circular-mapping mitochondrial genomes of *Calcarina hispida* and *Neorotalia gaimardi*. We also downloaded the available (meta)genomes of the foraminiferans *Reticulomyxa filosa* (*55*) and *Astrammina rara* (*56*) but could not identify mitochondrial genes.

### Retarian mitochondrial genomes have large intergenic regions

Since we found large intergenic regions in both foraminiferan and radiolarian mitochondrial genomes, we conducted searches for genes or gene fragments within these intergenic regions using blastx (v.2.11.0)(83), mfannot (https://github.com/BFL-lab/Mfannot), and hmmer (v3.3.2)(84). Twenty-four regions were identified as putatively homologous to genes typically encoded by rhizarian mitochondrial genomes (Figure 2 black lines). Eighteen of these are very similar to fragments of genes present elsewhere within the retarian mitochondrial genomes (*atp1*, *atp6*, *cob*, *nad4*, *nad5*, *nad7*, and *cox2*). The remaining six fragments are homologous to genes normally present in rhizarians, including *nad2* and *nad9* in radiolarians and *rps12* in a foram and a radiolarian. These fragments could represent pseudogenes, horizontally transferred DNA sequences (85, 86), or could reflect past genomic recombinations and rearrangements. In all mitochondrial contigs small ~50 bp stretches were nearly identical in many places only differing by a couple nucleotides. In the *Lithomelissa* sp. SAG, a large contig with similar read coverage was detected that contained these ~50 bp pseudo repeats but no mitochondrial genes or fragments (Figure 2C). Perhaps the missing mitochondrial rRNAs have diverged beyond recognition

### All three standard stop codons are likely recoded in radiolarian mitochondrial genomes

Deviations from the ancestral standard genetic code have evolved in numerous lineages (87, 88). In particular, lineages with extremely low GC content and limited opportunities for recombination (i.e., organellar genomes) exhibit genetic code changes more frequently (89, 90). One common trend of genetic code variability, and the easiest to detect, is when stop codons are reassigned as sense codons. The most common version of stop codon reassignment by far is the TGA stop codon being recoded to tryptophan (normally only encoded by TGG) (91). This change has occurred several times across mitochondrial genomes and in other bacterial lineages. The TAA and TAG stop codons can also be recoded. For example, in the mitochondrial genome of certain thraustochytrid stramenopiles, a new stop codon (TTA) (GenBank: AF288091.2) evolved and, in some species, both TAA and TAG were recoded to Tyrosine (normally encoded by TAT and TAC) (10). All three stop codons have been recoded in the nuclear genomes of the ciliate *Condylostoma magnum* (92) and the kinetoplastid genus *Blastocrithidia* (93, 94). In both cases, TGA = Tryptophan, and TAA and TAG = Glutamine (normally encoded only by CAA and CAG). For *Blastocrithidia*, authors showed that highly expressed genes have fewer TAA and TAG codons and speculate that changes in tRNA usage enable ribosomes to read through TAA and TAG sense codons in the middle of moderate and lowly expressed genes while TAA is still used as a termination codon at the end of transcripts (93, 94). Here, we identify a similar example in radiolarian mitochondria where all three stop codons are likely recoded to sense codons.

To determine the genetic code of radiolarian mitochondrial genomes, we aligned predicted proteins with mitochondria-encoded proteins from diverse protists. These alignments revealed that in-frame TGA and TAG codons occur at sites often occupied by tryptophan and tyrosine residues, respectively (Figure 3). Conversely, relatively few in-frame TAA codons are present in conserved domains. The majority of in-frame TAA codons occurred at locations for which there was no consensus amino acid in the alignment (Figure 3). In Acanthometra, only two genes contained in-frame TAAs (*nad5* (8xTAA) and *cox2* (2xTAA)). All eight *nad5* TAA codons were in the 3′ region, which appears to have diverged compared to the same region of other *nad5* genes. Similarly, the two *cox2* TAA codons were also in regions of the gene that are not highly conserved. In *Lithomelissa*, eight genes contained in-frame TAA codons. Like *Acanthometra*, *Lithomelissa* contained a few TAAs that aligned with conserved residues (one glutamine, one arginine, and the other tyrosine) in the middle of protein alignments (*nad7* and *cox3*).

**Figure 3.**
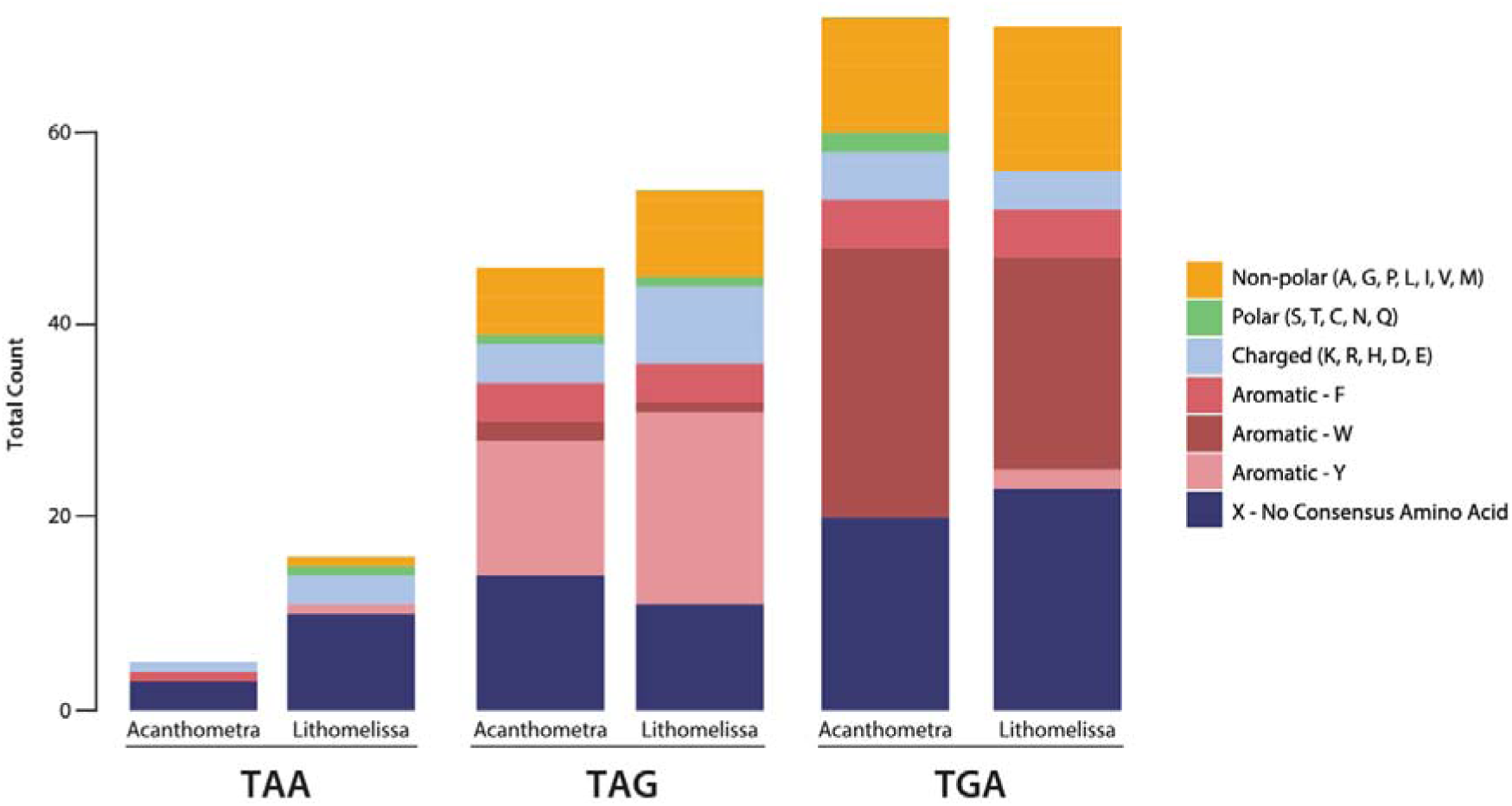
All three stop codons have been recoded to sense codons in radiolarian mitochondrial genomes. Proteins encoded in diverse mitochondrial genomes were aligned with their orthologues from radiolarians. Consensus (>50%) amino acids at sites containing internal radiolarian stop codons (TGA, TAG, TAA) were noted. All three stop codons have been recoded to sense codons in radiolarian mitochondrial genomes. Proteins encoded in diverse mitochondrial genomes were aligned with their orthologues from radiolarians. Consensus (>50%) amino acids at sites containing internal radiolarian stop codons (TGA, TAG, TAA) were noted. Amino acids were grouped based on their biochemical properties (non-polar, polar, charged, or aromatic). When a large proportion of sites are occupied by a particular amino acid, this suggests that the in-frame stop codon encodes that amino acid.

When assessing pairwise alignments of the radiolarian proteins (e.g., pairwise alignment of *Lithomelissa* and *Acanthometra cox1*), of the 29 in-frame TAA codons, nearly half (14xTAA) aligned with a tyrosine, phenylalanine, or tryptophan, and the majority (19xTAA) aligned with a hydrophobic residue. In addition, 25 of 27 radiolarian mitochondrial protein-coding genes had a TAA codon near where the end of the protein is predicted. Two *Acanthometra* proteins (*nad4L* and *cox2*) lacked stop codons and were contiguous with the open reading frames of *cob* and *nad1*, respectively. These data all suggest that a mechanism similar to the one proposed for the *Blastocrithidia* nuclear genome may be in place in radiolarian mitochondrial genomes. While TGA and TAG respectively encode tryptophan and tyrosine, TAA appears to have a dual role, likely encoding tyrosine in some proteins at a few locations but acting primarily as a stop codon. Curiously, several proteins lack any in-frame TAA codon. Perhaps, similar to the case in *Blastocrithidia*, the highest expressed mitochondrial proteins lack in-frame stop codons. The *atp9* gene is among the highest expressed and has no TAA or TAG present in either radiolarian. These data indicate that the mitochondrial genetic code in radiolarians has diverged from the ancestral code and has recoded all three stop codons to code for amino acids. While TGA and TAG are recoded to tryptophan and tyrosine, TAA codons sometimes encode for tyrosine but are the primary, and likely only, stop codon.

### Retarian mitochondrial genomes contain fragmented rRNA genes, divergent *atp6* genes, and split *nad* genes

In most eukaryotic lineages, mitochondrial genomes encode a combination of proteins involved in electron transport and ATP synthesis, ribosomal proteins, and a few auxiliary proteins involved in protein maturation or translocation (3). However, five major lineages (euglenids, retarians, chlorophycean algae, myzozoans, and animals - see Figure 4) have completely transferred all genes for mitoribosomal proteins to the nucleus, and two others are close behind (fungi encode only *rps3*, and glycomonads (Euglenozoa) encode at most *rps3* and *rps12*) (34, 77, 95–97). In all these lineages except animals and fungi, the EGT of mitoribosomal proteins has coincided with an extreme reduction or fragmentation of the mitochondrial rRNAs (Figure 4 dark blue circles) (98). Animal and fungal mitochondrial ribosomal RNAs are truncated, but not to the extent of other mitochondrial rRNAs that are extremely divergent and nearly undetectable.

**Figure 4.**
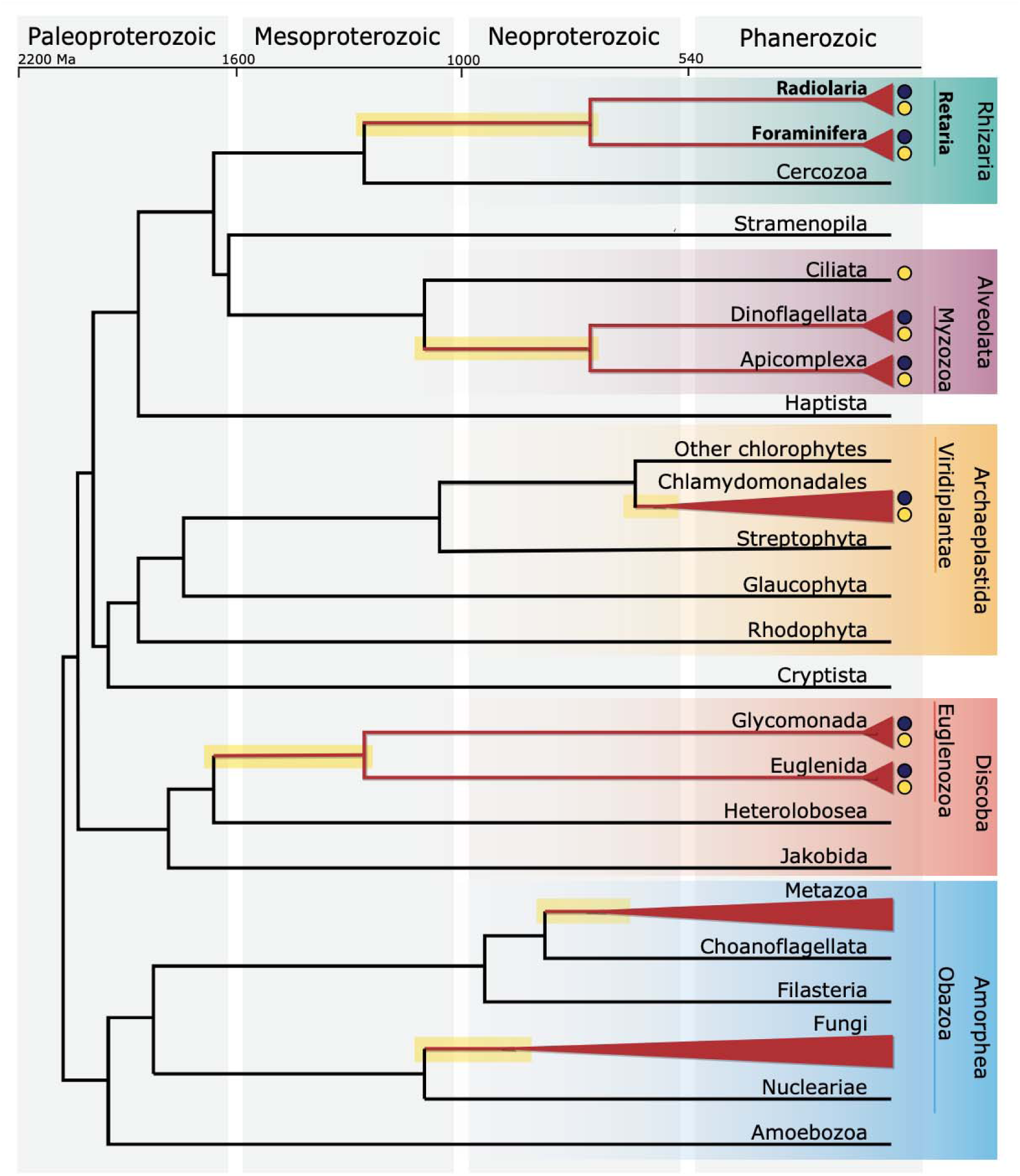
Divergent mitochondrial traits have persisted for hundreds of millions of years. Schematic phylogeny of extant eukaryotes with divergence times approximated based on (117). Clades highlighted in red have lost all mitoribosomal protein-coding genes from the mitochondrial genome. Dark blue circles indicate groups with short or fragmented mitochondrial rRNA genes. Yellow circles indicate groups with highly divergent mitochondrial atp6 genes. Lineages possessing these traits diverged in the mid-Neoproterozoic or earlier (emphasized with a branch highlighted in yellow).

In addition to highly divergent rRNA genes, euglenozoans, retarians, chlorophycean algae, and myzozoans possess extremely divergent *atp6* genes (Figure 4 yellow circles). Since a few TAA codons appear in the 5′ region of the *Lithomelissa atp6* gene, we decided to model the Atp6 protein using Alphafold2 to determine if the N-terminal extension is part of the protein or represents a non-coding upstream sequence. Alphafold2 modeled full-length subunit a into a structure that better resembles the classic subunit a (Figure S5). This suggests that the TAA codons are in part of the coding sequence of *Lithomelissa atp6*. Divergence of ATP synthase structure can have consequences for mitochondrial cristae morphologies (e.g., chlorophycean algae, euglenids, kinetoplastids, and apicomplexans all have unique cristae morphologies) (99–102). Since mitochondrial cristae architecture that departs from classic lamellar and tubular morphologies present in most other eukaryotes has also been reported for Foraminifera (103) and Radiolaria (104, 105), retarian ATP synthase structures represent excellent candidates for future investigation into the structural and functional diversity of this amazing protein complex (106).

Curiously, *nad1* is split into two parts in the Foraminifera *Calcarina hispida* and *Neorotalia gaimardi.* This suggests that some trans-splicing might be present in Retaria, similar to what has been reported for mitochondrial genes in other eukaryotes (107, 108). It is also possible that two peptides are separately expressed and merged into a functional protein, as has been found in *Chromera* plastids (109). Furthermore, we identified a conserved frameshift in *cox1* of all four analyzed species of the foraminifera order Miliolida, which suggests that a mechanism for stop codon read-through or post-transcriptional mRNA modification of this codon exists in this lineage. Manual insertion of a single ‘N’ into the miliolid sequences resulted in a continuous open reading frame (ORF), which, when translated, spans the entire length of the *cox1* protein sequence (110). Post-transcriptional insertion modifications have evolved in several protist lineages, including euglenids and diplonemids (95, 111). As the same pattern was found in all analyzed miliolid Foraminifera but not in any rotaliid species, we conclude that this is a unique feature of Miliolida mitochondria, which might be of interest in the future characterization of this group.

## Conclusions

Why do mitochondrial genomes vary so drastically across eukaryotes? Specifically, what triggers the wholesale transfer of mitochondrial ribosomal genes to the nucleus in so many lineages? There are several possible benefits to mitochondria-to-nucleus gene transfer (112), and given enough time, mitochondria-to-nucleus EGT is considered to be mathematically inevitable (113). Perhaps the diversity of mitochondrial genomes is simply a result of these evolutionary forces playing out over billions of years, with no functional cell biological consequences. However, this seems a somewhat unsatisfying answer given the diversity of mitoribosomal structures that have recently been solved (114–116).

In any case, retarian mitochondrial genomes represent a newly discovered ancient independent reduction in organellar gene content. The reduced gene complement of retarians displays more similarities to the mitochondrial genomes of euglenozoans, myzozoans, chlorophycean algae, animals, and fungi than it does to that of other rhizarians (Cercozoa). While several of these lineages may seem obscure and disparate, it is important to note that each lineage diverged in the mid-Neoproterozoic, or earlier (see (117) and Figure 4). These lineages therefore possess histories as deep and rich as animals and fungi, which are each traditionally considered to be independent ‘kingdoms’. Given the ancient divergence of forams and radiolarians, the strikingly reduced mitochondrial genomes of Retaria have persisted without substantial change for over 500 million years. The persistence of mitochondrial gene content over large time spans suggests that mitochondrial-to-nuclear gene transfer does not occur consistently but rather in relatively short macroevolutionary bursts. Further investigations into deeper branching taxa at nodes of apparent sudden mass EGT will clarify this notion. In sum, the retarian mitochondrial genomes presented here bridge a major gap in our understanding and provide the first glimpse into the mitochondria of this diverse group of ancient protists.

## Material and Methods

### Sample collection

#### Foraminifera samples

We analyzed 31 benthic Foraminifera cells (15 species) from the Spermonde Archipelago in Indonesia and from Coral Bay in Australia (see Table S1 for samples and locations). All specimens were stored in >90% ethanol after sampling and transferred to the Naturalis Biodiversity Centre laboratory for morphological species identification and molecular analyses. Specimens were sorted into morphotypes, identified and photographed using a ZeissDiscovery v12 stereo microscope (Oberkochen, Germany).

#### Radiolaria samples

Marine surface-water plankton samples were collected from the Pacific Ocean near the California coast (33.454219, −117.705215) by towing an 87 µ m-mesh plankton net from the back of a kayak on February 7, 2021, at 10:00 am. Bulk environmental plankton samples were immediately aliquoted into 15 mL falcon tubes and preserved with RNAlater. Plankton samples were stored on ice during transportation to the lab. Radiolarian cells were identified by morphology and imaged prior to single-cell isolation under an inverted microscope using a micropipette. Individual cells were washed four times in DNase and RNase-free water to remove extracellular material from each radiolarian. This process was repeated twice with new water each time before each cell was transferred to 4 uL of RNAlater and then stored at −20C before further processing.

### DNA extraction and sequencing

#### Foraminifera

Single Foraminifera specimens were dried in sterile 1.5-ml Eppendorf tubes and ground into a fine powder using a porcelain mortar and pestle. Total genomic DNA extraction was carried out using the QIAamp DNA Micro Kit (Qiagen; Hilden, Germany) as described in (110). After extraction, DNA quantification was conducted using the FragmentAnalyzer system (Agilent Technologies, Santa Clara, USA). Since extracted DNA was already fragmented to less than 500bp average length, no further fragmentation using ultrasonication or enzymes was conducted.

Shotgun metagenomic libraries were prepared using the New England Biolabs NEBNext Ultra II DNA Library Prep Kit (Ipswitch, USA) with the corresponding NEBNext Multiplex Oligos for Illumina, following the manufacturer’s protocol but reducing volumes by 50 percent. Final concentration and fragment size were checked on the Tapestation system (Agilent Technologies, Santa Clara, USA). All samples were equimolar pooled before sending for sequencing on the Illumina NovaSeq 6000 platform (2×150 bp read length) at Baseclear (Leiden, The Netherlands), targeting 5 million reads per sample.

#### Radiolaria

Single-cell DNA extractions were performed using the MasterPure DNA and RNA Purification Kit (Epicentre Biotechnologies) following the protocol as written, with the addition of a 30-minute incubation with a solution of lysis buffer and Proteinase K at 65C and 1000rpm. Purified total genomic DNA was eluted into 4uL TE Buffer and quantified using a Qubit HS dsDNA kit.

Genomic DNA from each cell was amplified using the Repli-G Advanced DNA Single-Cell kit and protocol (QIAGEN) for amplifying purified genomic DNA. Final concentration and fragment size were checked using the Tapestation and Qubit systems. An aliquot of each single-amplified genome containing a total of 500 ng DNA was provided to the ASU Genomics Facility for library preparation using KAPA Biosystem’s LTP library preparation kit before the samples were sequenced on the Illumina NovaSeq 6000 platform targeting 10 million 2×150 bp reads per sample.

### Bioinformatic analysis

#### Foraminifera

MultiQC (118) was used for the quality assessment of raw reads. Megahit (119) was used for the initial assembly of reads into contigs, which were loaded into Geneious Prime (v.2020) together with raw reads. Contigs were mapped against the mitochondrial genome of the rhizarian *Lotharella oceanica* deposited in GenBank (accession number NC_029731.1(77)) with up to 50% mismatch, a word length of 5, and up to 10% gaps (gap size 10) allowed. Since none of the assembled contigs could be mapped, raw reads and contigs were mapped against the *L. oceanica* reference with the same settings as above, and against Foraminifera mitochondrial COI barcode sequences published in (120). Regions with high coverage of mapped reads or with mapped contigs were manually inspected. In case mapped contigs did not represent a full mitochondrial genome (which was the case only for *Neorotalia gaimardi*) mapped reads were used as a reference for repeated mapping with the Geneious Prime mapper, with a minimum of 100 base pairs overlap, a maximum of 1% mismatch and no gaps allowed. Mapping was repeated until no further reads could be mapped. The resulting contigs were checked for open reading frames (ORFs) with mitochondrial translation table 4, which has previously been reported for protist mitochondrial genomes (10).

Contigs were submitted to the mfannot mitochondrial annotation web server of the University of Montréal (https://megasun.bch.umontreal.ca/cgi-bin/mfannot/mfannotInterface.pl). ORFs identified as cytochrome oxidase subunit 1 (*cox1*) were searched against the NCBI GenBank reference database (121) and the Foraminifera *cox1* database (120) using blastn to identify the *cox1* sequence stemming from putative symbionts and the putative foraminiferal *cox1*. Annotations were manually curated in Geneious Prime. ORFs that were not annotated by mfannot were translated to proteins, subject to transmembrane prediction with TMHMM (122) and searched against Pfam (123), UniProt (124), SwissProt (125) and Ensembl (126) databases using the hmmer web server (84) to check for potential matches with known mitochondrial genes. In case a complete mitochondrial genome could not be obtained, the putative foraminiferal mitochondrial genes were identified by mapping reads against the newly assembled *Calcarina hispida* and *Neorotalia gaimardi* mitochondrial genomes as described above.

To verify that foraminiferal mitochondrial genes could also be obtained from previously published datasets, we downloaded the *Globobulimina* (order Rotaliida) metagenome from the NCBI Sequence Read Archive (accession number: SRX3312059 (67)) and assembled the foraminiferal mitochondrial genes as described above. Furthermore, we downloaded the genomic contigs of the foraminiferans *Reticulomyxa filosa* (55) and *Astrammina rara* (*56*) and searched for mitochondrial genes as described above, though none could be found.

#### Radiolaria

MultiQC (118) was used to trim and filter raw fastq reads, which were then normalized with BBNorm (an addition of BBMap v.38.12). SAGs were assembled using SPAdes (v.3.15.2) (127). Normalized reads were mapped back to contigs with BBMap, and genome completeness was assessed with BUSCO (v.5.1.2) (128). Blobtools (v.1.0) (129) was used to visualize contigs with similar read coverage and GC content. Mitochondrial contigs were identified using *Andalucia godoyi* mitochondria-encoded proteins as queries in tblastn searches against radiolarian SAG assemblies. The mitochondrial contigs identified were manually stitched together by identifying regions of overlap >50bp between contigs with similar read coverages.

Putative mitochondrial contigs were submitted to the mfannot mitochondrial annotation (https://megasun.bch.umontreal.ca/apps/mfannot/) web server. Because mfannot did not identify full-length rRNA genes within our mitochondrial genomes, nhmmer (130) was used to search each genome for rRNA genes using manually curated rRNA databases. Fragments of mitochondrial genes were also identified by searching intergenic regions and open reading frames that were not annotated by mfannot against a manually curated database of mitochondrial protein sequences with representatives from all protist genera with a sequenced mitochondrial genome in NCBI GenBank using blastx. Intergenic regions and ORFs with at least four hits from the same gene were considered significant enough for annotation on the mitochondrial genome maps. Annotations were added manually using Geneious Prime.

### Stop Codon Analysis

Amino acid multiple sequence alignments were used to assess the locations within a mitochondrial gene at which radiolarians have a stop codon. Alignments were generated with MUSCLE (131) using radiolarian genes identified by mfannot and genes from every available protistan genus on GenBank. If more than one mitochondrial genome existed for a genus on Genbank, then the most recent two mitochondrial genomes from different species were chosen as representatives for that genus. The total number of stop codons present within each mitochondrial gene from the two radiolarian mitochondrial genomes was visually counted. The 50% consensus amino acid identities at locations for which a radiolarian mitochondrial gene had an in-frame stop codon were also tallied to assess which amino acids radiolarian stop codons are potentially coding for instead. Radiolarian stop codons that occurred at locations where the consensus alignment contained a gap or where the majority of genes within the alignment were not present (the very beginnings and ends of the alignment) were not counted. Pairwise alignments of radiolarian mitochondria-encoded proteins were performed using MUSCLE and inspected manually.

### Phylogenetic analysis of Foraminifera and Radiolaria

Twelve mitochondrial protein-coding genes (*cox1*, *cox2*, *cox3*, *cob*, *nad3*, *nad4*, *nad4L*, *nad5*, *nad7*, *atp1*, *atp6*, *atp9*) were aligned with MAFFT (v7.450) (132). The split *nad1* gene was excluded from phylogenetic analyses. Aligned protein sequences per gene were manually trimmed to the same length, and stop codons were removed. All analyzed genes were manually concatenated. Gaps in the alignment were manually removed, resulting in an alignment of 2,137 amino acids. A phylogenetic tree was calculated using the IQ-TREE web server (133) with the JTT+F+G4 model and 1,000 iterations of Ultrafast Bootstrap(133). We visualized the resulting tree using FigTree (v1.4.4) (Rambout, available online: https://github.com/rambaut/figtree/).

## Data availability

Raw reads are available in the NCBI Sequence Read Archive (SRA): BioProject PRJNA743004.

The full mitochondrial genomes of *Calcarina hispida* and *Neorotalia gaimardi* have been deposited to NCBI GenBank (accession numbers OP965949 & OP965950). The radiolarian genome assemblies, multiple sequence alignments, and predicted mitochondrial gene sequences have been deposited to Figshare: doi: 10.6084/m9.figshare.16734961. Assemblies can be searched using BLAST on a SequenceServer (134) at https://evocellbio.com/SAGdb/macher_et_al/.

## Acknowledgements

This material is based upon work supported by the National Science Foundation under Grant No. DBI-2119963 (JGW). We are grateful to T. Coots for assistance with computational analyses.

## Supplementary Figures

**Figure S1.**
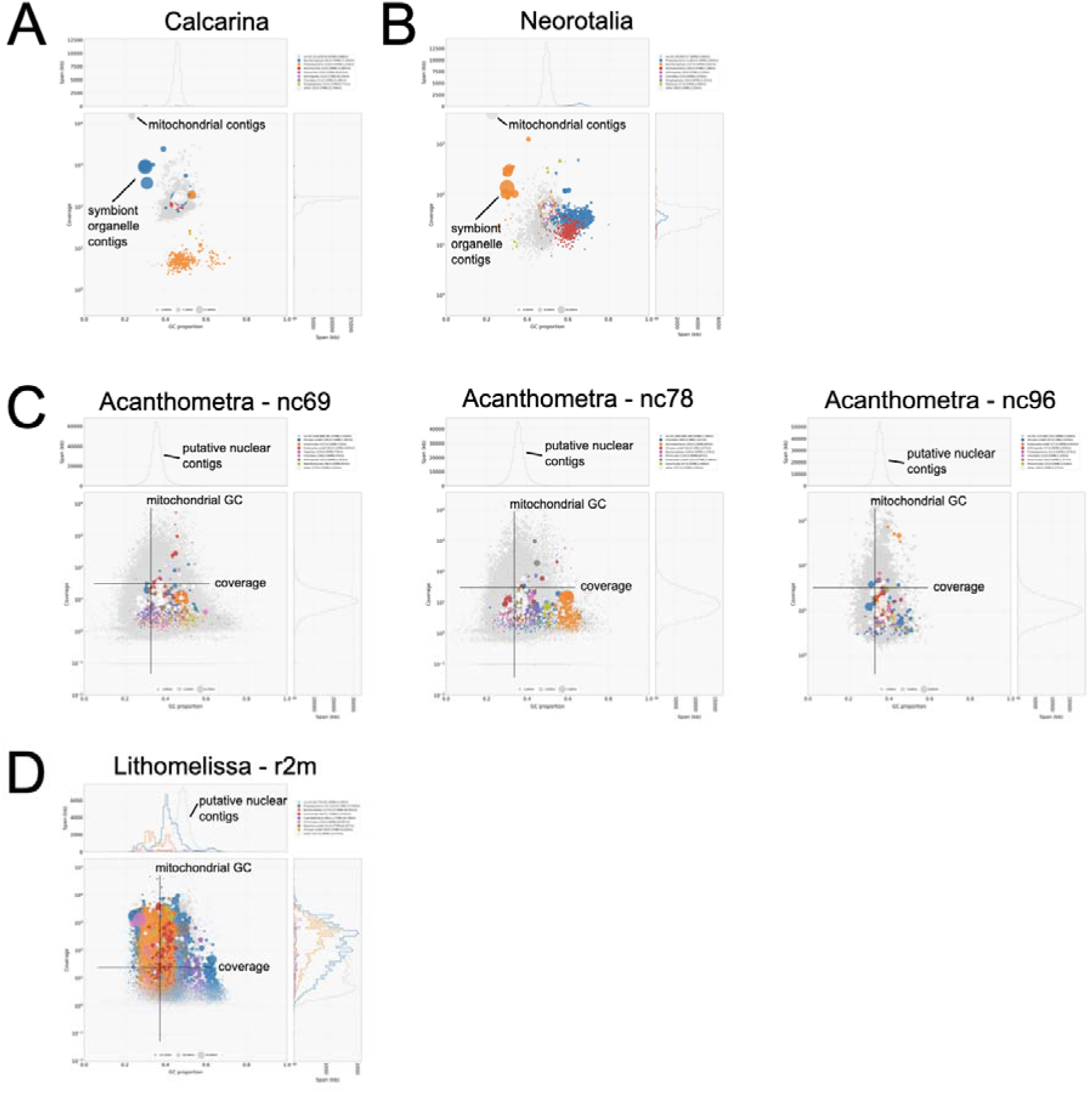
BlobTools plots of retarian SAGs. (A) *Calcarina* blobplot. (B) *Neorotalia* blobplot. (C) *Acanthometra* blobplots. (D) *Lithomelissa* blobplot. Mitochondrial contigs and symbiont organelle contigs are highlighted in foram blobs (A and B). The average GC content and coverage of mitochondrial contigs are indicated in radiolarian blobs (C and D).

**Figure S2.**
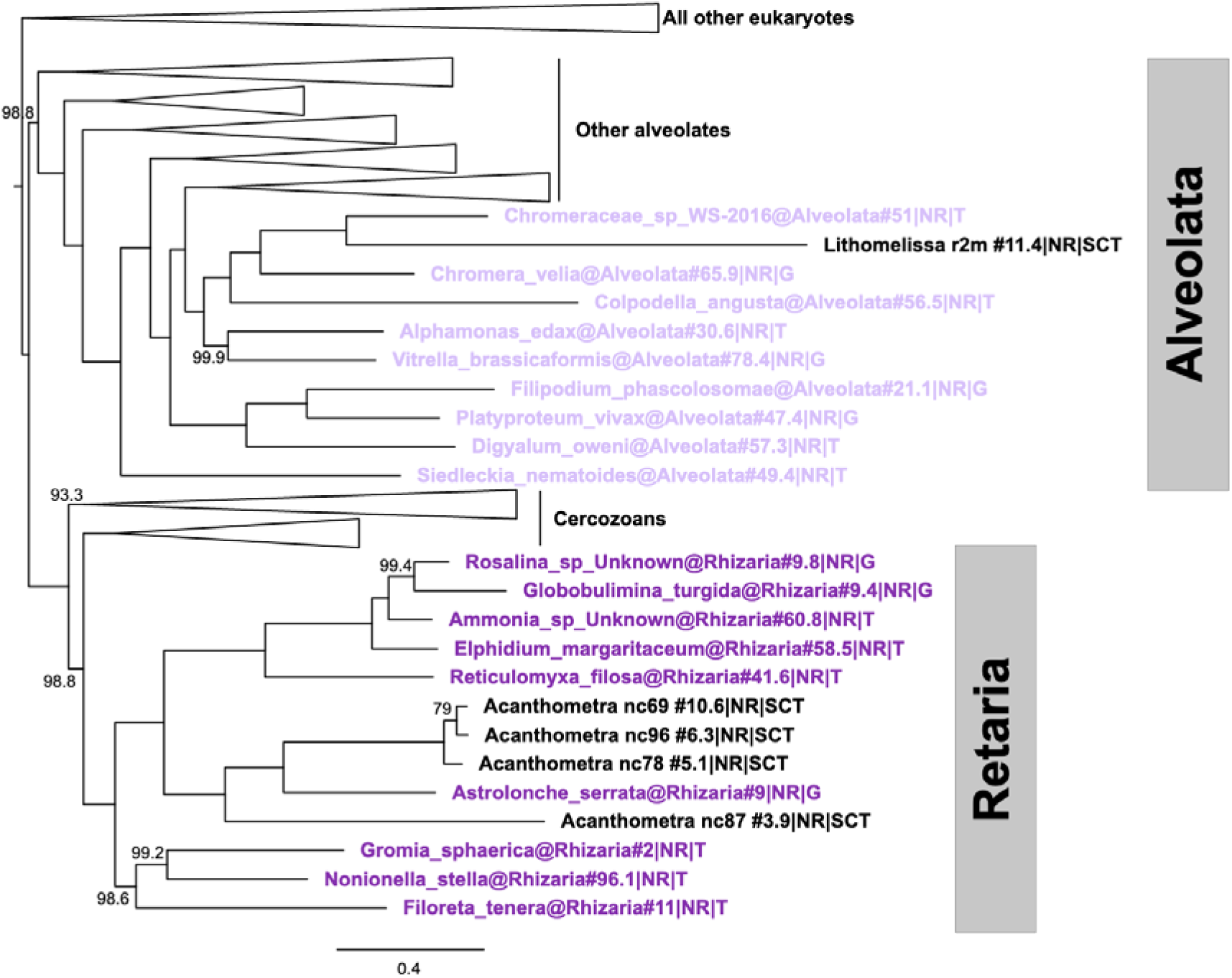
Phylogenetic placement of radiolarian SAGs. BUSCO proteins retrieved from radiolarian SAGs were added to alignments previously generated (EukProt (71)). *Acanthometra* and *Amphibelone* SAGs were placed with the lone radiolarian in the dataset. The *Lithomelissa* SAG was placed within alveolates, likely due to contamination.

**Figure S3.**
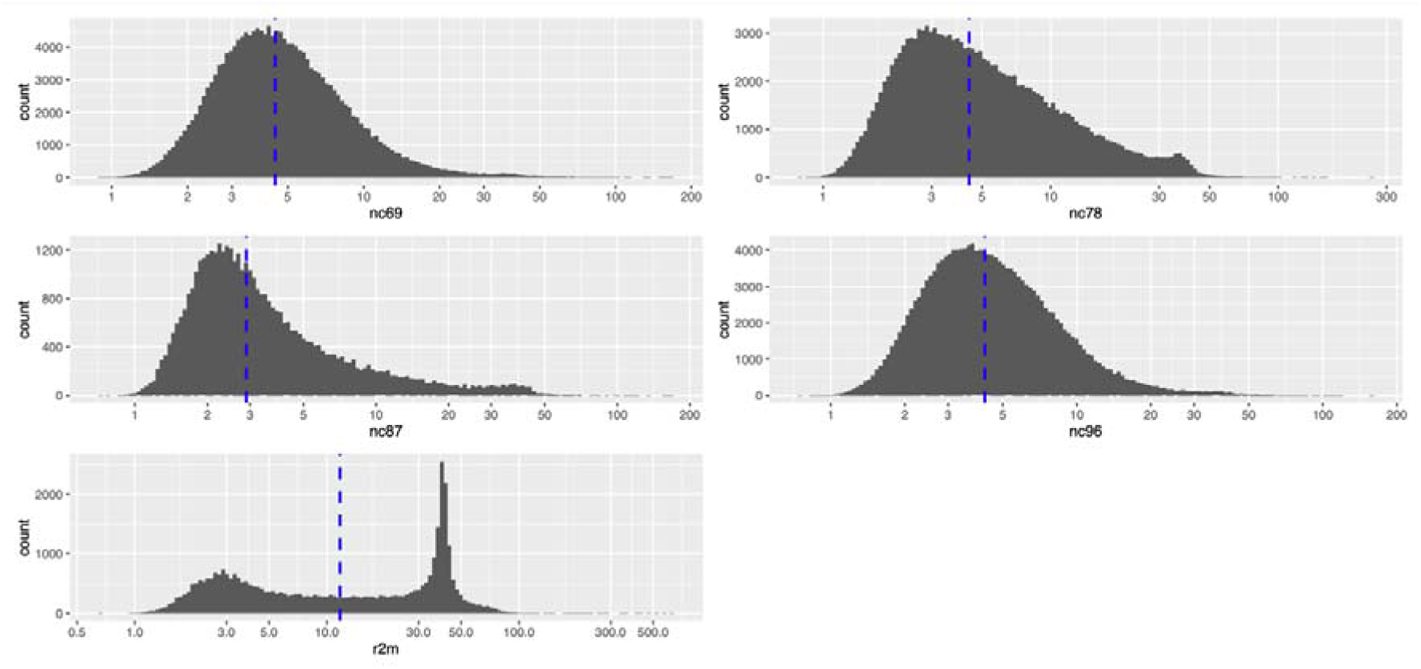
Coverage plots for radiolarian SAGs. The number of contigs versus read coverage was plotted. The median coverage is indicated in blue.

**Figure S4.**
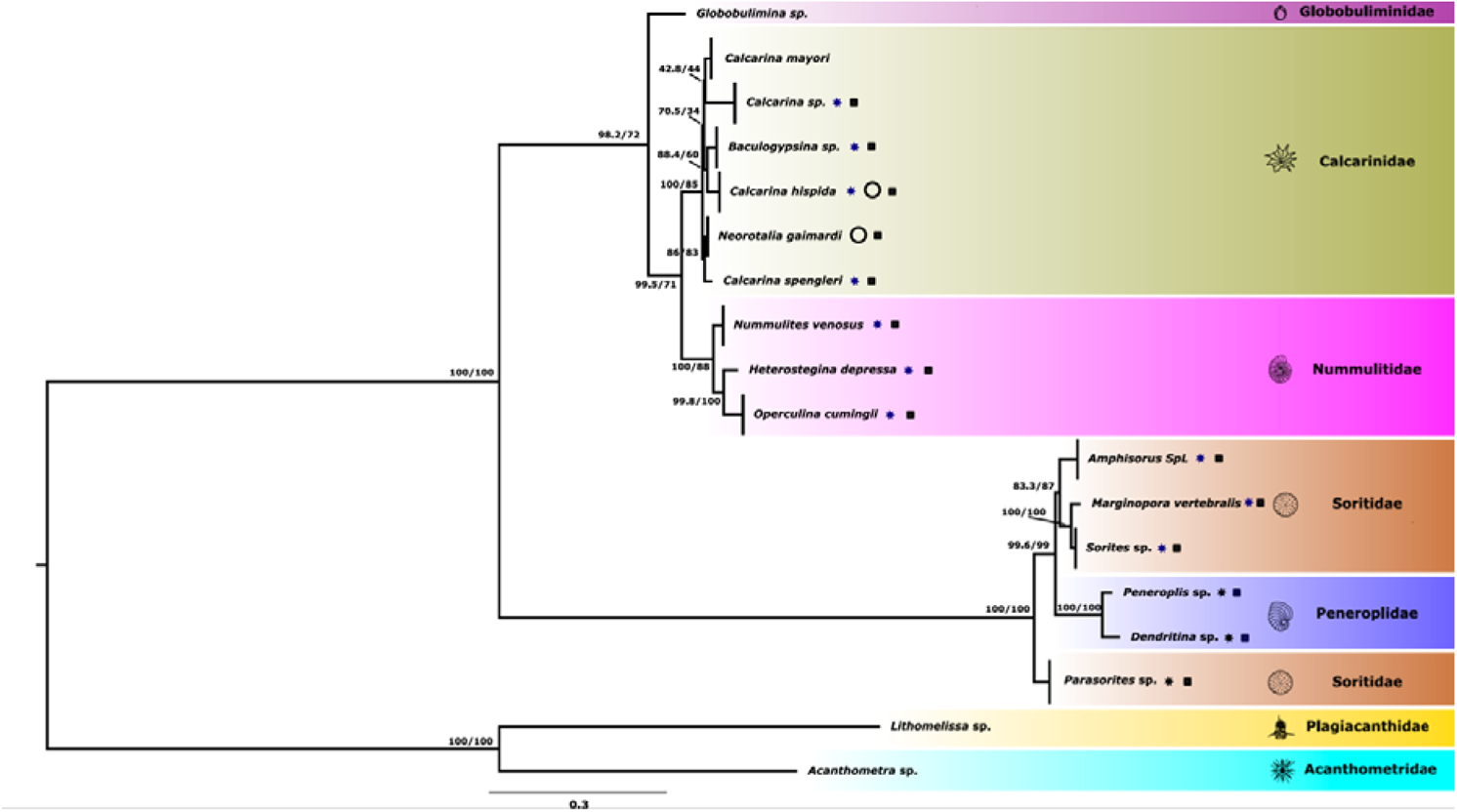
Foraminifera phylogenetic reconstruction using concatenated mitochondrial proteins confirms SSU trees. The phylogeny of Foraminifera was inferred using Radiolaria as an outgroup. Twelve mitochondrial proteins from nine Foraminifera and two Radiolaria species were concatenated (2471 amino acids) into a data matrix. The tree was calculated using IQ-Tree with the JTT+F+G4 model and 1000 iterations (82). Numbers at nodes indicate bootstrap values (Maximum likelihood) and posterior probabilities (Bayesian inference). Branches below genus level are collapsed. We visualised the resulting tree using FigTree (v1.4.4) (Rambout, available online: https://github.com/rambaut/figtree/). Stars behind species names indicate recovery of 18S rRNA for the species; a circle indicates the recovery of a circular mitochondrial genome, and a black square indicates the recovery of symbiont 28S (for dinoflagellate-bearing forams) or symbiont rbcl (for diatom-bearing forams)

